# Identifying candidate genetic determinants associated with invasion in pneumococcal lineages

**DOI:** 10.64898/2026.10.27.762133

**Authors:** Maiju Pesonen, Oliver Lorenz, Mignon du Plessis, Nicole Wolter, Shabir A. Madhi, Susan A. Nzenze, Anne von Gottberg, Nicholas J Croucher, Keith P. Klugman, The Global Pneumococcal Sequencing Consortium, Jukka Corander, Stephen D. Bentley, Stephanie W Lo, Rebecca A. Gladstone

## Abstract

Serotype invasiveness estimates help identify high-risk serotypes and interpret disease replacement following pneumococcal conjugate vaccination. Because serotype and lineage are strongly associated, their independent contributions must be separated to identify unexpectedly invasive lineages and investigate underlying mechanisms.

Using temporally matched carriage and invasive-disease isolates from children aged <7 years in South Africa and the USA, we fitted a logistic mixed-effects model with Global Pneumococcal Sequence Cluster (GPSC) as a random effect and serotype and country as fixed effects. We conducted a species-wide genome-wide association study (GWAS) of invasion and compared the prevalence of significant unitigs in GPSC1 and GPSC21 to identify invasion-associated variants differentiating them. We additionally examined capsule-locus variation that could contribute to their contrasting invasiveness.

After serotype and country adjustment, GPSC1 and GPSC21, which both express 19F and share an immediate common ancestor, ranked fifth and 149th of 150 lineages; odds multipliers were 1.51 (95% conditional interval 1.06–2.15) and 0.61 (0.41–0.93), respectively. Of 1,569 invasion-associated unitigs from the species-wide GWAS, 272 differentiated GPSC1 from GPSC21. Among these, the strongest signal mapped to *hk11* (*desK*) and was associated with GPSC1; *bgaA* was the highest-ranked known antibody-binding target. Within GPSC1 CC271 had additional variants in the ATP-dependent Clp protease ATP-binding subunit ClpX compared to the CC420 subclade. Capsule-protein folds were conserved with local effects indistinguishable from prediction uncertainty.

Separating serotype and lineage effects can identify unexpectedly invasive lineages and prioritize candidate serotype-independent determinants, vaccine antigens and capsule variation for experimental investigation.

**Data Summary:** Code implementing the mixed-effects invasiveness model is available at https://github.com/rgladstone/pneumo-lineage-invasion. Deidentified isolate metadata, sequence accessions, analysis inclusion and supporting data underlying the figures and tables are available in the Supplementary Tables.

**Impact Statement:** Among pneumococcal isolates collected from children younger than 7 years, pneumococci with the same capsule type differed in their capacity to cause invasive disease. Separating the effects of capsule type and genetic lineage makes it possible to identify unexpectedly invasive lineages and investigate the reasons for their increased disease potential. Applying this approach highlighted candidate proteins that may help explain invasiveness and could be explored as vaccine antigens with relevance beyond individual lineages. Several differences were found in and around the capsule-producing region; predicted protein structures were conserved, although subtle effects on capsule composition or expression cannot be ruled out. This work therefore provides a way to uncover both non-capsular factors and overlooked capsule variation associated with disease, helping to prioritize mechanisms, vaccine candidates and structures for experimental investigation.

## Introduction

The association of pneumococcal serotypes with invasive pneumococcal disease (IPD) varies markedly. Serotypes 1, 8 and 12F are regularly reported with the highest associations, whilst other serotypes are more associated with carriage; including 6C and 11A, along with non-typeable isolates that are rarely isolated from invasive sites [1, 2]. Young children are generally considered the principal reservoir for pneumococcal carriage and transmission, although carriage prevalence, acquisition and transmission are strongly age-dependent [3, 4]. Estimates of invasiveness for pneumococcal serotypes are fundamental to understanding the differing disease potential of pneumococci. However, serotype and lineage are non-independently distributed: some serotypes are concentrated within few lineages, whereas others, including 19F, occur across many. Consequently, conventional odds ratios cannot separate serotype from lineage effects where they are strongly linked.

Capsular serotype is established as a major determinant of pneumococcal invasiveness [5], but numerous non-capsular virulence factors have experimentally supported roles in pathogenesis. For example, the pore-forming toxin pneumolysin damages host cells and modulates innate immune responses, whereas surface proteins such as pneumococcal surface protein A (PspA) and pneumococcal surface protein C (PspC) facilitate adhesion to host cells and evasion of complement-mediated immunity [6]. The widespread conservation of pneumococcal surface proteins across the species, together with their accessibility to the immune system, has made them attractive candidates for serotype-independent vaccines [7, 8]. We previously reported that the heritability of invasion from the bacterial genetics rather than host factors was 0.57, and that serotype only explained half of that heritability [9, 10]. This suggests that serotype-independent factors will have a polygenic influence on invasiveness that remains understudied. Furthermore, the persistence of vaccine serotypes such as 3 and 19A, [11] together with the continuing restructuring of pneumococcal populations and the consequent expansion of vaccine valency, underscores the need for alternative vaccine targets. Investigating associations between pneumococcal genetic variation and disease phenotypes, for example, through genome wide association studies (GWAS), provides a means of prioritising known virulence factors and identifying novel candidates for functional validation and vaccine development [10, 12]. Again, clonality results in variation, for example, allelic variants and accessory gene content, which is lineage specific, that require innovative solutions to tease apart associations with invasion and lineage effects [13, 14].

We aimed to first identify highly invasive lineages and subsequently identify the putative genetic determinants via their association with both the disease phenotype itself and particular lineages. We also endeavored to identify further variation in the capsular polysaccharide synthesis (*cps*) locus that could impact capsular phenotype between isolates with the same serotype designation [15, 16]. Global Pneumococcal Sequence Clusters GPSC1 and GPSC21 have been indicated by multiple approaches to differ strikingly in invasiveness when expressing the 19F serotype [9, 17], giving an opportunity to discover virulence factors with a measurable impact on disease progression. Moreover, these lineages share an immediate common ancestor in the species-wide phylogeny reducing the total pool of differentiating genetic factors [18]. We therefore aimed to further refine lineage and serotype invasiveness quantification using an interpretable logistic mixed model, and to identify genetic determinants that might explain the difference between lineages with invasiveness left unexplained by current serotype designations.

## Results

### The invasive potential of serotypes and lineages

Serotype fixed-effect estimates varied substantially, and residual between-lineage variation remained after adjustment for serotype and country (Figure 1). Among children younger than 7 years, lineages expressing the same serotype differed in estimated invasiveness, indicating lineage-associated variation beyond that explained by serotype and country. The lineage-specific odds multipliers estimated by the model for the three highest-ranked lineages were numerically similar to the model-estimated odds ratios for serotypes 6C through to 11A relative to non-typeable isolates. Strikingly, the estimates for the relative invasiveness of two 19F expressing lineages, GPSC1 that predominantly expresses 19F and 19A, and GPSC21 which is almost exclusively 19F, were at opposing ends of the distribution of random effects across all tested lineages after controlling for serotype. These two lineages are known to share an immediate common ancestor, with the individual lineages previously estimated to be more than 150 years old [18]. GPSC1 is globally distributed, though a strong geographical population structure exists, where isolates from the USA cluster and are predominantly 19A whilst South African isolates expressing 19F cluster distinctly in the phylogeny (Figure 2A). Country contributed to the estimated lineage effects but did not account for the elevated invasiveness associated with GPSC1. In a sensitivity analysis, our adjustment for country reduced the GPSC1 lineage log-odds effect by 23%, corresponding to a reduction in its lineage odds multiplier from 1.71 (95% conditional interval 1.19–2.45) to 1.51 (1.06–2.15). The USA had different case-to-carriage sampling proportions, and inclusion of country reduced the estimated between-lineage variance by 47%. Nevertheless, the GPSC1 effect was retained in a South-Africa-only model (odds multiplier 1.89, 95% conditional interval 1.10–3.25). Consistent with this result, a direct comparison restricted to South African serotype 19F isolates showed higher disease odds for GPSC1 than GPSC21 (OR 4.25, 95% CI 1.73–10.54; Fisher’s p=0.00064).

**Figure 1.**
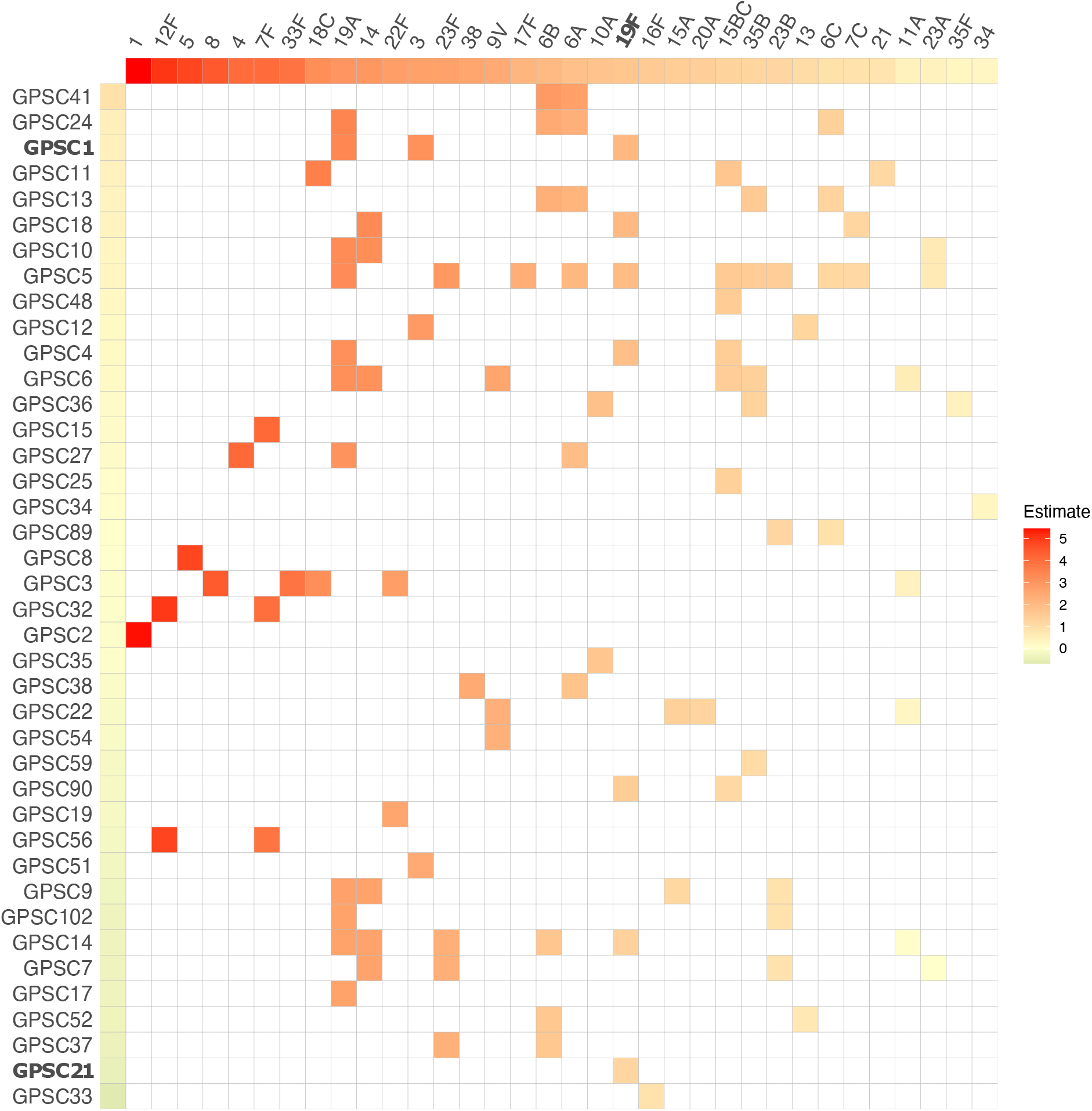
The estimated marginal and combined invasive potential of different serotypes and lineages. The colours represent regression coefficients on a logit scale (log-odds) estimated from a generalised mixed model with clinical manifestation (infection/carriage) as a binary outcome, serotypes as a fixed effect and lineages as random effects. Lineages are denoted as Global Pneumococcal Sequencing Clusters (GPSCs). Red tones are associated with higher, and green tones with lower estimates of invasive potential. The serotype reference category was classified by non-typeable (NT) phenotype, or NT genotype when phenotype was unknown. Serotype fixed effects and GPSC random effects were estimated simultaneously. Columns are ordered by fixed-effect estimates and rows independently by adjusted lineage random effects, both from highest to lowest; the ordering therefore does not reflect the combined value of individual serotype–GPSC cells.

**Figure 2.**
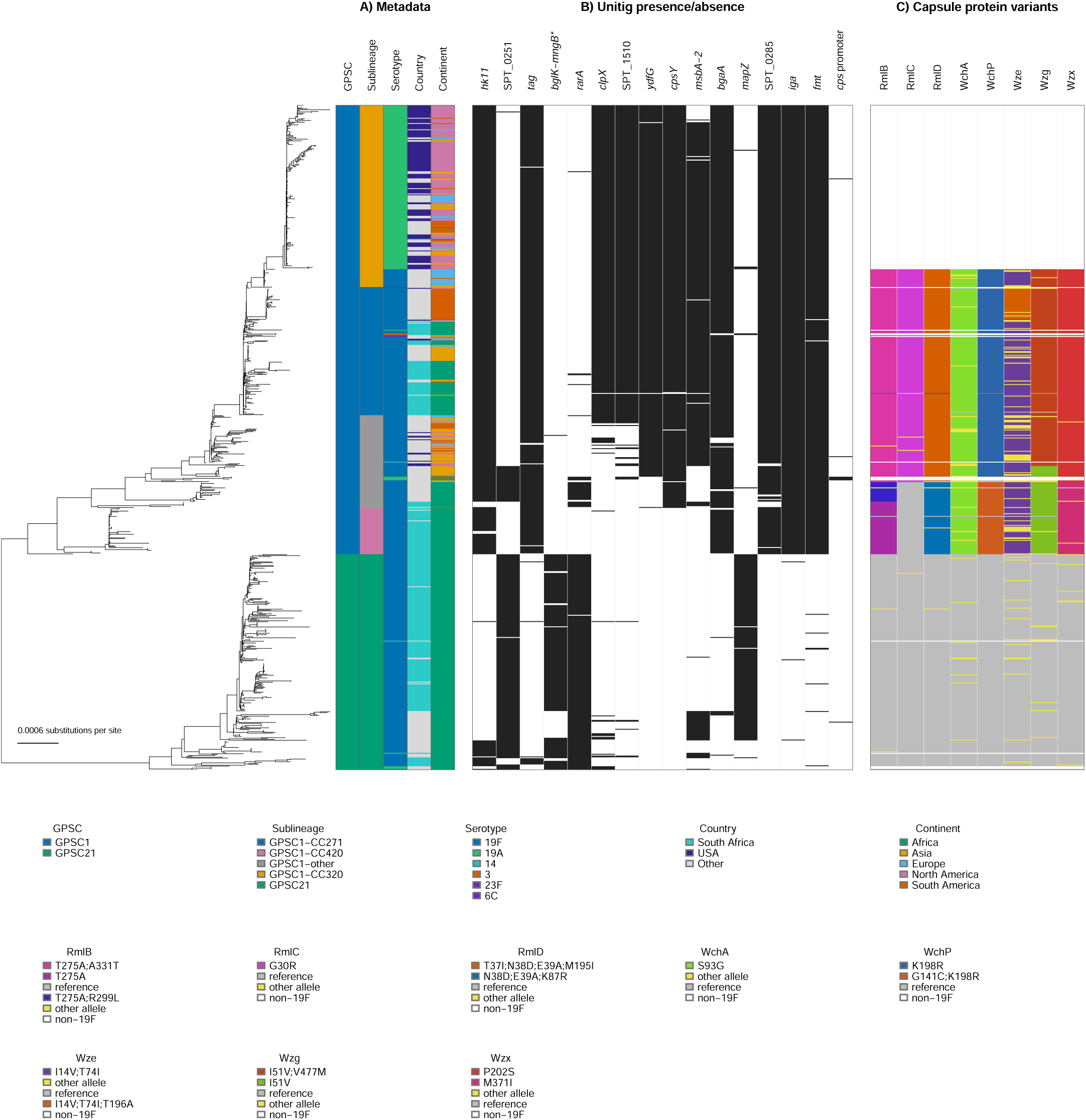
Phylogenetic structure of GPSC1 and GPSC21 and features. Midpoint-rooted maximum-likelihood phylogeny of 746 GPSC1 and GPSC21 isolates. A) metadata associated with phylogenetic structure B) the unitig presence (black) or absence (white) of 16 selected lineage-associated unitigs and the cps promoter unitig; locus tags are shown where no gene designation was available, and the asterisk denotes the intergenic *bglK– mngB* unitig. C) selected cps gene amino-acid profiles for RmlB, RmlC, RmlD, WchA, WchP, Wze, Wzg and Wzx relative to the GPSC21 reference. Proteins were selected to show the principal GPSC1–GPSC21 differences and secondary variation within GPSC1. Common annotated profiles are shown individually; substitutions separated by semicolons occur in the same protein profile. Grey indicates identity to the GPSC21 reference, yellow denotes other called alleles pooled outside the displayed common profiles, and white denotes non-19F isolates. An interactive version of this figure is available at https://microreact.org/project/oSRNi5uzwyMzQabEKjmdYx-gpsc1-gpsc21

Furthermore, there are two discrete clusters of South African isolates in GPSC1, corresponding to clonal complexes CC271 (56.3% IPD) and CC420 (38.0% IPD). Conversely, GPSC21 was only observed in Sub-Saharan Africa, and only 16% of these South African isolates were from IPD. Lineage definitions can be applied at different resolutions, and these proportions of invasive-disease isolates suggested an invasiveness gradient from the GPSC1 CC271 sub-lineage (9/16; 56.3%) through CC420 (8/21; 38.1%) to GPSC21 (18/110; 16.4%). Although point estimates suggest a difference in invasiveness in the GPSC1 sub-lineages, the confidence intervals overlapped and did not support a definitive ordering of their invasiveness. The isolates included in these analyses are denoted in Supplementary T1. The invasiveness model effect estimates, and sensitivity analysis are included in Supplementary T2 and Supplementary T3 respectively.

### Unitigs associated with invasion status and lineage invasiveness

We performed a species-wide genome-wide association study (GWAS) comparing 855 invasive-disease isolates with 1,089 carriage isolates from South African children aged <7 years, using a linear mixed model that accounted for population structure. This analysis identified 1,569 unitigs significantly associated with isolate status (invasive disease versus carriage). Using the same input unitigs, we additionally performed a fixed-effects GWAS of invasive-disease versus carriage status to assess lineage effects, which identified one significant unitig assigned to GPSC1 and none to GPSC21. Although GPSC1 and GPSC21 share an immediate common ancestor, 47,666 of the 903,113 unitigs tested among serotype 19F isolates (5.3%) differentiated GPSC1 from GPSC21, irrespective of GWAS significance, illustrating the large lineage-associated genetic variation space from which invasion-associated candidates had to be prioritised. Of these, 23,554 were present in ≥95% of GPSC1 and <5% of GPSC21 isolates, whereas 24,112 showed the reciprocal distribution.

We therefore filtered the hits from the linear mixed-model genome-wide association study (LMM-GWAS) for unitigs that also differentiated GPSC1 and GPSC21. This reduced the broad association signal to a focused set of unitigs that were both associated with invasive disease and characteristic of these lineages with substantial unexplained differences in invasive potential. Of the 1,569 significant unitigs, 272 (17.3%) differentiated the two lineages and mapped to 82 distinct gene or locus contexts; these unitigs were not restricted to the focal lineages. Of these, 141 were characteristic of GPSC1, and all had β > 0, indicating a positive association with invasive disease. The remaining 131 were characteristic of GPSC21, and all had β < 0, indicating a negative association with invasive disease (Supplementary Figure 1).

Thus, intersecting lineage differentiation with disease association substantially narrowed the extensive background of lineage-associated genomic variation to a smaller and more biologically interpretable set of candidate loci. The differentiating significant unitig with the smallest p-value had a positive β value, was associated with GPSC1 and was ranked 36^th^ in the full GWAS hits (Table 1A). This unitig was observed in a core gene encoding the sensor histidine kinase of pneumococcal two-component regulatory system 11, *hk11* (annotated *desK)*, which was the 6th gene ranked by the number of significant hits in the invasion GWAS. A total of 19 unitigs in this gene differentiated the two lineages, with 89% of them having a positive β value for invasion.

**Table 1.** Gene context for selected species-wide invasion-associated unitigs differentiating (A) GPSC1 and GPSC21 and (B) GPSC1-CC271 from GPSC1-CC420 and GPSC21. Gene names and products use the best-supported pneumococcal annotation. Locus tags refer to Taiwan19F-14 (GCF_000019025_1, GPSC1); a dash denotes an unassigned gene name. Lineage differentiation was assessed among the 166 South African serotype 19F isolates belonging to GPSC1 (n=48) or GPSC21 (n=118) that were included in the species-wide GWAS. Differentiating unitigs were defined as present in ≥95% of one group and <5% of the comparison group, ordered by species-wide invasion-GWAS likelihood-ratio-test p-value and collapsed by gene/locus context. Panel A shows the five highest-ranked GPSC1/GPSC21 unitig locus contexts. Panel B shows the five highest-ranked locus contexts represented by unitigs present in ≥95% of CC271 and <5% of both CC420 and GPSC21. Rank is the position among the 1,569 significant species-wide GWAS unitigs. The n/N columns summarize all differentiating or significant unitigs assigned to each locus context, not only the representative unitig. Positive and negative β coefficients indicate associations with invasive disease and carriage, respectively.

**A. GPSC1 versus GPSC21.**
| Gene; locus tag | Description | Best species-wide GWAS rank (among lineage-differentiating unitigs) | Lineage-differentiating/ all significant unitigs, n/N | Positive $\beta$ among GPSC1-associated unitigs, n/N | Positive $\beta$ among GPSC21-associated unitigs, n/N | Positive $\beta$ among all significant unitigs, n/N |
| --- | --- | --- | --- | --- | --- | --- |
| *hk11*<br>SPT_1982 | TCS11 sensor histidine kinase | 36 | 19/28 | 17/17 | 0/2 | 17/28 |
| —<br>SPT_0251 | Short hypothetical; immediately upstream of a GNAT-family acetyltransferase | 47 | 5/10 | 0/0 | 0/5 | 0/10 |
| *tag*<br>SPT_0217 | DNA-3-methyladenine glycosylase I | 51 | 8/10 | 7/7 | 0/1 | 7/10 |
| —<br>SPT_2155-SPT_2156 | Intergenic; between the ROK-family β-glucoside kinase BglK and α-mannosidase MngB | 52 | 2/2 | 0/0 | 0/2 | 0/2 |
| *rarA*<br>SPT_1717 | Replication-associated recombination protein A | 92 | 8/11 | 4/4 | 0/4 | 6/11 |

**B. GPSC1-CC271-associated unitigs distinguishing CC271 from both GPSC1-CC420 and GPSC21**
| Gene; locus tag | Description | Best species-wide GWAS rank (CC271-specific pattern) | Lineage-differentiating/ all significant unitigs, n/N | Positive $\beta$ among CC271-associated unitigs, n/N | Positive $\beta$ among CC420-associated unitigs, n/N | Positive $\beta$ among all significant unitigs, n/N |
| --- | --- | --- | --- | --- | --- | --- |
| *clpX*<br>SPT_1509 | ATP-dependent Clp protease ATP-binding subunit ClpX | 31 | 60/117 | 14/14 | 0/46 | 30/117 |
| —<br>SPT_1510 | Hypothetical; between dihydrofolate reductase and the ATP-dependent Clp protease ATP-binding subunit ClpX | 135 | 13/21 | 6/6 | 0/7 | 6/21 |
| *ydfG*<br>SPT_1858 | Short-chain dehydrogenase/reductase-family oxidoreductase | 173 | 1/2 | 1/1 | 0/0 | 2/2 |
| *cpsY*<br>SPT_0700 | MtaR-like LysR-family transcriptional regulator | 242 | 12/12 | 2/2 | 0/10 | 2/12 |
| *msbA-2*<br>SPT_0917 | ABC transporter ATP-binding/permease protein | 252 | 14/22 | 6/7 | 2/7 | 14/22 |

We further filtered the LMM-GWAS hits for differential unitig presence between the GPSC1 sublineages CC271 and CC420, reducing the signal to invasion-associated unitigs that differentiated sublineages with a potential gradient in invasive potential. Despite their closer phylogenetic relationship, CC271 and CC420 were differentiated by nearly as many unitigs as GPSC1 and GPSC21 (46,949 versus 47,666, respectively). However, the CC271–CC420 unitigs showed substantially greater co-segregation, suggesting that the raw count reflects extensive linked or lineage-structured variation rather than an equivalent number of independent genomic differences. Of the 1,569 significant unitigs, 224 differentiated the two GPSC1 sublineages and mapped to 56 distinct gene or locus contexts. The direction of association was consistent with the higher invasiveness point estimate for CC271 than for CC420: 75 of 87 (86.2%) CC271-associated unitigs were positively associated with invasive disease, whereas 123 of 137 (89.8%) CC420-associated unitigs were negatively associated with invasive disease. For the final Table 1B prioritisation, we added GPSC21 as a third comparator to exclude CC271-associated unitigs that were also common in GPSC21 and therefore did not follow the observed gradient in disease association. Table 1B presents the five highest-ranked locus contexts represented by unitigs present in ≥95% of CC271 isolates and <5% of both CC420 and GPSC21 isolates. Of the 87 CC271-associated unitigs, 76 met this additional GPSC21 criterion and 75/76 had positive β coefficients. The highest-ranked qualifying unitig mapped to *clpX*, was present in 22/22 CC271, 0/26 CC420 and 1/118 GPSC21 isolates, had β = 1.24 and ranked 31^st^ in the full species-wide GWAS. All 14 CC271-associated *clpX*-mapping unitigs had positive β coefficients.

Finally, we determined whether the 272 unitigs that differentiated GPSC1 from GPSC21 were observed in known immunogenic targets previously identified in a pangenome-wide immunological screen [19]. Only 5 of the 82 distinct gene/locus contexts corresponded to known immunogenic targets (Table 2). Among genes encoding known antibody-binding targets, *bgaA* - a cell-wall-anchored β-galactosidase - was the highest ranked. All six GPSC1-associated lineage-differentiating *bgaA* unitigs had positive β values. Of further interest, three GPSC1-associated unitigs mapped to *iga* (*zmpA*), and all had positive β values.

**Table 2.**
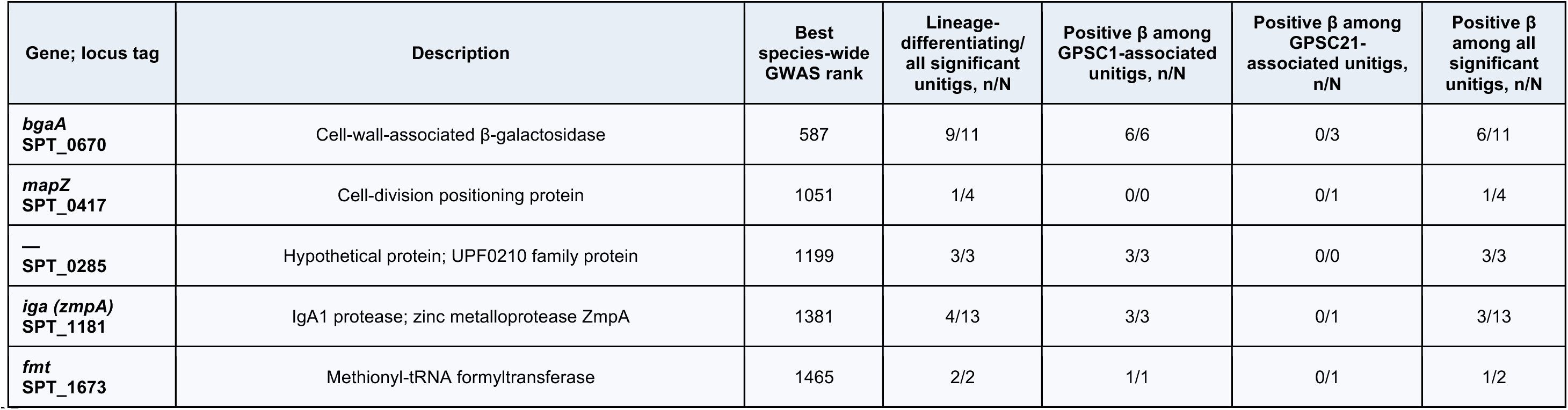
Gene context for all five genes encoding previously reported antibody-binding targets among the species-wide invasion-associated unitigs that differentiated GPSC1 and GPSC21. Gene names and protein descriptions use the best-supported pneumococcal annotation; locus tags refer to Taiwan19F-14 (GCF_000019025_1, GPSC1). A dash indicates that no gene name is assigned. Lineage differentiation was assessed among the 166 South African serotype 19F isolates belonging to GPSC1 (n=48) or GPSC21 (n=118) that were included in the species-wide GWAS. Significant species-wide GWAS unitigs were considered lineage-differentiating when present in ≥95% of one lineage and <5% of the other. All five genes encoding previously reported antibody-binding targets that met this criterion are shown. Rank gives the position of the best lineage-differentiating unitig for each gene among the 1,569 significant species-wide GWAS unitigs, ordered by likelihood-ratio-test p-value; identical p-values shared a rank. Positive β values indicate association with invasive disease.

| Gene; locus tag | Description | Best species-wide GWAS rank | Lineage-differentiating/<br>all significant unitigs, n/N | Positive $\beta$ among GPSC1-associated unitigs, n/N | Positive $\beta$ among GPSC21-associated unitigs, n/N | Positive $\beta$ among all significant unitigs, n/N |
| --- | --- | --- | --- | --- | --- | --- |
| *bgaA*<br>SPT_0670 | Cell-wall-associated β-galactosidase | 587 | 9/11 | 6/6 | 0/3 | 6/11 |
| *mapZ*<br>SPT_0417 | Cell-division positioning protein | 1051 | 1/4 | 0/0 | 0/1 | 1/4 |
| —<br>SPT_0285 | Hypothetical protein; UPF0210 family protein | 1199 | 3/3 | 3/3 | 0/0 | 3/3 |
| *iga (zmpA)*<br>SPT_1181 | IgA1 protease; zinc metalloprotease ZmpA | 1381 | 4/13 | 3/3 | 0/1 | 3/13 |
| *fmt*<br>SPT_1673 | Methionyl-tRNA formyltransferase | 1465 | 2/2 | 1/1 | 0/1 | 1/2 |

Among South African GWAS isolates belonging to GPSCs other than GPSC1 or GPSC21, one significant invasion-associated unitig mapped across the *cps* core promoter and contained a C- to-T substitution immediately downstream of the predicted −10 element (TATAATC→TATAATT). This T-containing unitig occurred in 139/1,944 South African isolates (7.2%) and was positively associated with invasion (β=2.83). A C-containing unitig from the same promoter region occurred in 1,461/1,944 isolates (75.2%) and had a negative effect estimate (β=−0.813). These unitig-presence patterns were not mutually exclusive: 33 isolates contained only the T-containing pattern, 1,355 only the C-containing pattern, 106 both and 450 neither. Consequently, these unitig calls could not be used to assign a definitive C or T promoter genotype to every isolate. The T-containing unitig occurred predominantly in GPSC2 (serotype 1), GPSC18 (serotype 14), GPSC125 (serotype 9N) and GPSC137 (serotype 9N). It was also detected in 6/504 GPSC1 and 1/242 GPSC21 isolates from three other African countries, China and Brazil, but not among the South African GPSC1 or GPSC21 isolates. The GWAS results for all 1,569 significant unitigs are detailed in Supplementary T4. The distributions of the selected unitigs within GPSC1 and GPSC21 are displayed in Figure 2B and detailed in Supplementary T5.

### Capsule polysaccharide locus variation

No significant species-wide invasion-associated GWAS unitigs mapped to the GPSC1 or GPSC21 serotype 19F *cps* coding sequences. However, a *cps* effect confined to the GPSC1– GPSC21 contrast would be lineage-specific and might not be captured as a species-wide association after correction for population structure. We therefore examined their *cps* loci directly to identify variation that might plausibly alter capsule expression or sugar structure.

Gene order was conserved across the GPSC1 and GPSC21 19F *cps* loci. The core promoter upstream of the capsule regulatory gene *wzg* was also conserved for these lineages; upstream differences were limited principally to two promoter-spacer SNPs and distinct mobile-element architectures observed in the GPSC1 and GPSC21 hybrid genomes (Figure 3).

**Figure 3.**
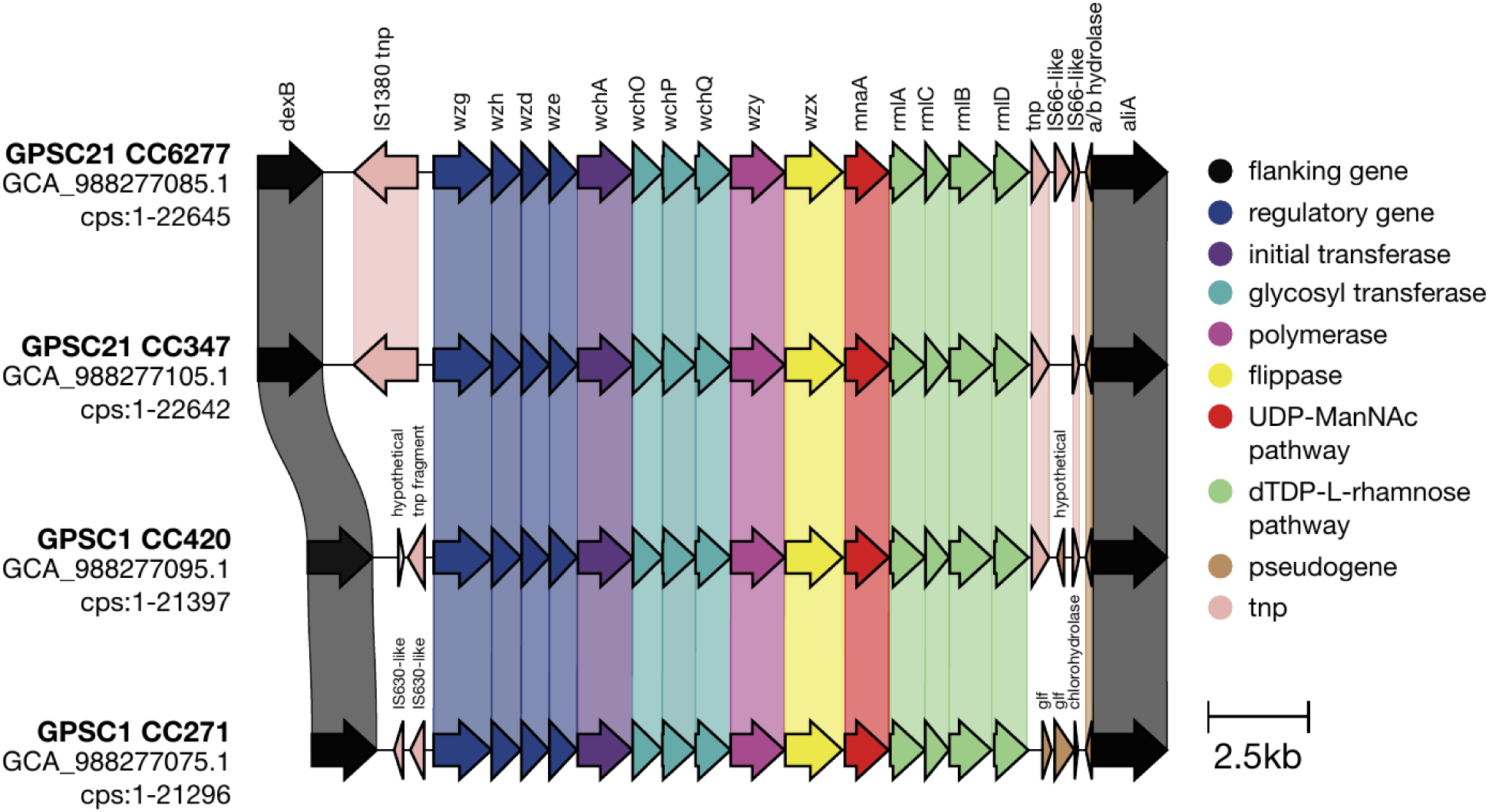
Comparison of GPSC1 and GPSC21 capsular polysaccharide biosynthesis gene clusters. Gene organization is shown for representative isolates from GPSC21 (CC6277 and CC347) and GPSC1 (CC420 and CC271). Arrows indicate coding sequence position and orientation; gene names are shown above the upper cluster, with variable or additional genes labelled in the corresponding clusters. Colors denote functional categories, including transposases (tnp). Shaded links connect homologous genes between adjacent clusters, with sequence-similarity >0.96.

At the protein sequence level, a WchP K198R substitution distinguished GPSC1 from GPSC21 among South African serotype-19F isolates. R198 was present in all GPSC1-CC271 isolates (69/69) and most GPSC1-CC420 isolates (47/48) but was absent from GPSC21 (0/180).

Despite this strong lineage association, the K198R-centred regions remained structurally similar between the representative alleles (Cα RMSD, 0.407–0.469 Å). The dominant *rmlD* and *rmlB* alleles contained stepwise radical amino-acid substitutions—replacements between residues with substantially different physicochemical properties—that mirrored the invasiveness gradient of GPSC1-CC271 -> GPSC1-CC420 -> GPSC21. AlphaFold predicted highly similar global and substitution-centred backbone structures for these alleles (global RMSD, 0.092–0.170 Å; local RMSD, 0.051–0.101 Å), finding no evidence of major structural changes, although effects on electrostatics, side-chain interactions or enzyme kinetics cannot be excluded.

RmlC G30R was fixed in GPSC1-CC271 (69/69) but absent from GPSC1-CC420 (0/48) and GPSC21 (0/180), while WchP G141C was present in 46/48 GPSC1-CC420 isolates but absent from GPSC1-CC271 (0/69) and GPSC21 (0/180). Comparisons of the top-ranked AlphaFold models showed local RMSDs of 1.115 Å for RmlC G30R and 1.984–2.311 Å for WchP G141C, while their global folds remained conserved. However, these local RMSDs were smaller than the corresponding mean local PAE values of 1.61–1.91 Å and 4.38–5.28 Å, respectively.

Consequently, no robust local structural alteration distinguishable from model uncertainty was detected for either substitution. These substitutions were retained as hypotheses for functional testing based on their lineage distributions and predicted biochemical roles, rather than as evidence of altered protein structure.

The remaining amino-acid variants showed no compelling predicted backbone alterations. Together, these findings provide little evidence that major capsule remodeling accounts for the greater invasiveness of GPSC1, although RmlC G30R and WchP G141C remain candidates for biochemical and functional testing. The lineage distributions of selected cps protein variants are displayed in Figure 2C. The CPS protein allele profiles and promoter/upstream-region variation are detailed in Supplementary T6 with full structural comparisons and confidence metrics are provided in the Supplementary T7.

## Discussion

Among carriage and invasive-disease isolates from children younger than 7 years, we found a marked difference between closely related lineages expressing the same serotype. GPSC1 and GPSC21 share an immediate common ancestor and both express 19F, yet after serotype and country adjustment they ranked fifth and 149th of 150 GPSCs, with estimated lineage odds multipliers of 1.51 (95% conditional interval 1.06–2.15) and 0.61 (0.41–0.93), respectively—a 2.5-fold contrast between point estimates. This quantifies substantial lineage-associated variation and complements previous estimates of pneumococcal disease progression [17]. In the South African species-wide GWAS, 1,569 unitigs were associated with whether isolates came from invasive disease or carriage; 272 also differentiated GPSC1 and GPSC21, mapping to 82 gene or locus contexts. Among these, the highest-ranked signal mapped to *hk11* (β = 1.08; Benjamini–Hochberg-adjusted p = 1.14 × 10⁻⁸), while *bgaA* was the highest-ranked known antibody-binding target (β = 0.93; BH-adjusted p = 5.67 × 10⁻⁶). GPSC1-associated unitigs also mapped to *iga,* with β values of 0.93–0.96 and BH-adjusted p = 2.17 × 10⁻⁵ for the highest-ranked unitig. Within GPSC1, 224 GWAS-significant unitigs differentiated CC271 from CC420, with 86.2% and 89.8% of the respective sublineage-associated unitigs showing directions consistent with the observed disease gradient. The highest-ranked CC271-specific candidate, also rare in CC420 and GPSC21, mapped to *clpX* (β = 1.24; BH-adjusted p = 7.53 × 10⁻⁹).

GPSC21 was concentrated in South Africa, with too few isolates observed in other countries. The genomic comparison therefore necessarily relied on South African isolates, limiting assessment of whether the candidate associations extend to other populations. Further work should test whether these associations correspond to specific phenotypic or biochemical differences, to explore how the candidate variants may contribute to invasiveness. No unitigs meeting the species-wide disease-association threshold mapped to the focal lineages’ 19F *cps* coding sequences, and structural predictions provided no convincing evidence of differences in overall folds or local backbone structures of the capsule proteins examined. However, effects on side-chain interactions, electrostatics, conformational dynamics, oligomerisation or enzymatic activity cannot be assessed reliably from static monomer predictions and remain possible modifiers.

Filtering for differences between GPSC1 and GPSC21 will miss determinants that do not differentiate the pair.

The candidates identified in this study have potential links between invasiveness and bacterial signalling, stress responses and host interactions. Within the two-component regulatory system 11 (TCS11), *hk11* encodes the sensor histidine kinase. Histidine kinases detect environmental signals and transduce them through phosphorylation of cognate response regulators, which in turn modulate downstream cellular processes, often including gene expression [20].

Experimental work has shown that TCS11 regulates pneumococcal colony-phase variation, favouring the opaque phenotype, while both components of TCS11—the response regulator RR11 and histidine kinase HK11 —were identified as putative transmission factors in a genome-wide transposon-sequencing (Tn-seq) screen [20, 21]. Allelic variation in *hk11* could plausibly alter signal sensing, kinase activity or phosphotransfer to RR11, potentially changing TCS11-regulated phenotypes relevant to colonisation, transmission and invasive disease [22]. In pneumococci, ClpP negatively regulates competence and contributes to stress adaptation, while ClpX was proposed as a possible partnering ATPase [23]. Variation in *clpX* could therefore influence invasiveness through altered competence or stress responses. Of the pneumococcal proteins previously demonstrated to be antibody binding targets, BgaA stood out. BgaA is a cell-wall-anchored β-galactosidase involved in host-glycan processing and epithelial adhesion, making its allelic variation both a plausible contributor to host interaction and an accessible antigenic target [19, 24]. Variation in the immunoglobulin A1 (IgA1) protease gene *iga* (also known as *zmpA*) provided a second interesting candidate: all three GPSC1-associated unitigs had positive β values, and its encoded surface-exposed zinc metalloprotease cleaves human IgA1, providing a direct potential link to immune evasion [25].

Whilst structural predictions provided no convincing evidence that the lineage-associated capsule-protein substitutions altered protein backbone structure, these substitutions remain potential modifiers of the amount, composition or linkage structure of the capsule produced. RmlC participates in dTDP-L-rhamnose biosynthesis, while WchP is predicted to transfer L-rhamnose-1-phosphate to ManNAc during assembly of the serotype 19F repeat unit [26, 27]. Given the diversity of pneumococcal *cps* loci [15, 27], and the substantial work required to confirm novel serotypes, differences in invasiveness between lineages expressing the same serotype provide a basis for prioritising capsule variants for functional investigation.

Closely related pneumococcal lineages expressing the same serotype differed substantially in invasiveness, again showing that serotype alone does not capture all variation in disease association. By intersecting species-wide disease associations with genomic differences between sister lineages with contrasting levels of invasiveness, we prioritised candidates relevant both across the pneumococcal population and within the closely related genetic backgrounds. Together, these analyses connect differences in invasiveness to specific genetic hypotheses, helping direct functional investigation towards variation that distinguishes lineages expressing the same serotype.

## Methods

### Invasiveness estimates

To estimate serotype and lineage invasiveness, we used published serotype and Global Pneumococcal Sequence Cluster (GPSC) data [9, 18] from carriage and nationwide invasive-disease collections in children younger than 7 years, excluding individuals known to be HIV-positive. Carriage and disease isolates were matched by sampling year within each country: 2009–2013 in South Africa and 2001, 2004, 2007 and 2009 in the USA. For USA isolates with no genomic data but known ST and serotype (n=217) the GPSC was inferred from the ST via the lookup table available at https://www.pneumogen.net/gps/#/gpsc#gpsc-st-lookup-table.

Within each country, isolates were randomly subsampled to maintain an approximately constant carriage-to-disease ratio across sampling years. The carriage-to-disease ratio—approximately 1.74:1 in South Africa and 0.76:1 in the USA—was set by the year with the fewest isolates of the category being subsampled relative to the category retained in full, thereby minimising exclusions. Separate country-specific ratios accommodated the opposing sampling balances of the South African and USA collections: carriage predominated in South Africa, whereas disease predominated in the USA (South Africa: n = 1,714, 36.5% invasive disease; USA: n = 801, 57.0% invasive disease). We fitted a two-level logistic mixed-effects model in R version 4.5.1 using the glmer function in the lme4 package (version 1.1-35.5) [28]. To reduce sparse-data problems and support model convergence, we further excluded serotypes represented by ≤2 carriage or disease isolates or ≤15 isolates overall. These criteria excluded 26 serotypes comprising 114 isolates, leaving 2,401 isolates for analysis. Lineage was included as a random intercept, with serotype and country included as fixed effects; non-typeable isolates were used as the reference category for serotype. The non-typeable reference category included isolates with NT phenotypes, or NT genotypes when phenotype was unknown, and comprised 37 isolates: 34 from carriage and three from invasive disease. As a sensitivity analysis for geographical confounding, we compared lineage effects estimated with and without country as a fixed effect and refitted the model using South African isolates only.

### GWAS and lineage comparisons

We performed a genome-wide association study on South African genomic data from children <7 years old, excluding known positive HIV individuals (n=1944). Combining countries could generate associations driven by country-specific sampling, serotype composition or phylogenetic structure rather than invasiveness. Furthermore, GPSC21 was concentrated in South Africa, whereas GPSC1 exhibited marked country-associated population structure: the US population was predominantly serotype 19A, while the South African GPSC1 population was predominantly serotype 19F. Unitigs were counted from the South African assemblies using unitig-caller v1.3.0 [29]. An initial GWAS to detect lineage effects was performed in pyseer v1.3.10 [14]. Lineage effects were assessed using the bugwas-style ‘--lineag’ workflow, with GPSC assignments supplied as custom lineage clusters. Serotype and ten multidimensional-scaling components derived from the genomic distance matrix were included as fixed effects to control for serotype and population structure, respectively; unitigs were assigned to the single GPSC with which their presence–absence pattern was most strongly associated. A second GWAS to identify variants associated with invasion was performed using a linear mixed model with a binary phenotype for the cases (IPD) and controls (carriage) and serotype as a covariate in pyseer v1.3.10 [14]. Population structure was corrected for using the kinship matrix. We determined a significance threshold using the number of unique unitig patterns 0.05/490,378 to be 1.01962E-07. The 1,569 significant unitigs were filtered to identify those that discriminate GPSC1 from GPSC21, with a presence in ≥95% in one and <5% of the other, and vice versa for all 166 GPSC1 and GPSC21 South African serotype 19F isolates in the GWAS: 22 CC271, 26 CC420 and 118 GPSC21. For Table 1B, CC271-associated unitigs were additionally required to be present in ≥95% of CC271 isolates and <5% of both CC420 and GPSC21 isolates; representative unitigs were ordered by full species-wide GWAS LRT p-value and collapsed by gene/locus context. All isolates belonging to GPSC1 and GPSC21 were annotated with ggCaller v1.3.4 [30], and the annotated de Bruijn graph (DBG) was queried with the filtered significant unitigs to provide gene information. Unitigs were additionally mapped to hybrid Taiwan19F-14 (GCF_000019025_1, GPSC1), GPSC1 and GPSC21 *in-house* hybrid assemblies, using Bowtie2 v2.3.5 [31–33]. Genes with significant hits were cross-referenced with those that had previously been shown to be immunogenic [19].

All Illumina data from GPSC1 and GPSC21 collected as part of the GPS project [9] were used to create a single phylogeny of these sister lineages for visualization purposes using IQ-TREE v2.0.3 [34] on an alignment from split k-mer analysis version 2 v0.2.0 (SKA2) [35] and recombination removed with Gubbins v3.4.3 [36], a fully interactive version was created with Microreact [37]. Published dated trees were used to establish the lineage root dates [18].

### Capsule polysaccharide locus variation

We compared the serotype 19F capsule loci of GPSC1 and GPSC21 at three levels: gene organisation, protein-allele and promoter/upstream sequence variation, and predicted structures of selected protein alleles. To create reference assemblies for representative strains in GPSC1 and GPSC21 we selected four isolates belonging to GPSC1-ST271, GPSC1-ST420, GPSC21-ST347 and GPSC21-ST6277 that had previously been sequenced with Illumina and resequenced them with Oxford Nanopore Technology. The fastq sets were provided to Hybracter v0.10.1 [38]. We annotated with Bakta v1.10.3 database v5 [39], extracted the *cps* locus between *dexB* and *aliA* and compared the gene synteny with Clinker v0.0.32 [40].

Protein-allele distributions were compared across the 297 South African serotype 19F isolates in GPSC1 and GPSC21. Capsule-gene coding sequences were translated and compared with the predominant GPSC21 protein allele for each gene to identify amino-acid substitutions. Allele and substitution frequencies were summarised within each group to identify differences between lineages and variation within GPSC1. Substitutions were classified by changes in residue physicochemical properties, including hydrophobicity, polarity and charge. For isolates with available upstream sequences, aligned regions preceding *wzg* were compared for nucleotide substitutions and insertions/deletions within the annotated core promoter, spacer and repeat unit of pneumococcus (RUP) regions. Upstream mobile-element organisation was compared separately in the four hybrid reference assemblies. Protein-allele profiles and promoter/upstream variants are provided in Supplementary Table T6.

Representative CPS protein alleles from GPSC1-CC271, GPSC1-CC420 and GPSC21 were modelled individually with AlphaFold 3 using AlphaFold Server (accessed July 2026) [41]. The top-ranked model_0.cif structure for each allele was superposed in ChimeraX v1.11.1 using MatchMaker [42, 43]. Global similarity was evaluated using pruned and all-pair Cα RMSDs, while local comparisons used substitution-centred residue windows. Model confidence was assessed using pTM, site-specific and window-mean pLDDT, and local PAE values [41, 44]. The structural comparisons were treated as exploratory.

### Supplementary data

Supplementary Figure 1 – GPSC1 and GPSC21 differentiating unitigs Manhattan plot Supplementary Tables – Accessions, metadata, results tables

### Funding information

The presented work received funding from a Pfizer supported Robert Austrian Research Award in Pneumococcal Vaccinology from the 12^th^ International Symposium on Pneumococci and Pneumococcal Disease (ISPPD-12, R.A.G). This work was additionally supported, in part, from Wellcome Trust (grant reference 206194; S.D.B.) through the Global Pneumococcal Sequencing project and by the Bill & Melinda Gates Foundation (under Investment ID INV-003570 to S.A.M., A.v.G., M.d.P., S.D.B. and S.A.N.). The conclusions and opinions expressed in this work are those of the author(s) alone and shall not be attributed to the Foundation. Under the grant conditions of the Foundation, a Creative Commons Attribution 4.0 License has already been assigned to the Author Accepted Manuscript version that might arise from this submission. Please note works submitted as a preprint have not undergone a peer review process. The funders had no role in study design, data collection and interpretation, or the decision to submit the work for publication.

### Author contributions

Conceptualization: R.A.G.; Data curation: M.P., O.L., S.A.N., S.W.L., R.A.G.; Formal analysis: M.P., O.L., R.A.G.; Funding acquisition: S.W.L., S.D.B., R.A.G.; Investigation: M.P., O.L., R.A.G.; Methodology: M.P., O.L., J.C., N.J.C., R.A.G.; Project administration: S.W.L., R.A.G.;

Resources: M.d.P., N.W., S.A.M., A.v.G., K.P.K., L.M., J.C., S.D.B., The Global Pneumococcal

Sequencing Consortium.; Software: M.P., O.L., R.A.G.; Supervision: S.W.L., J.C., S.D.B.,

R.A.G.; Validation: M.P., R.A.G.; Visualization: M.P., R.A.G.; Writing – original draft: R.A.G.; Writing – review & editing: M.P., O.L., M.d.P., N.W., S.A.M., S.A.N., A.v.G., J.C., N.J.C., K.P.K., L.M., S.D.B., S.W.L., R.A.G.

### Ethical approval and consent to participate

This study used published de-identified bacterial genomic data and metadata.

## Acknowledgements

We would like to thank all Global Pneumococcal Sequencing project partners and the ISPPD board for supporting this work. OpenAI Codex desktop application (OpenAI; accessed July 2026) was used under author direction to assist with verification of analyses and manuscript formatting. All code and outputs were independently reviewed and validated by the authors; the tool was not used to generate primary data, make analytical decisions or scientific conclusions.

---

**Supplementary Figure 1.**
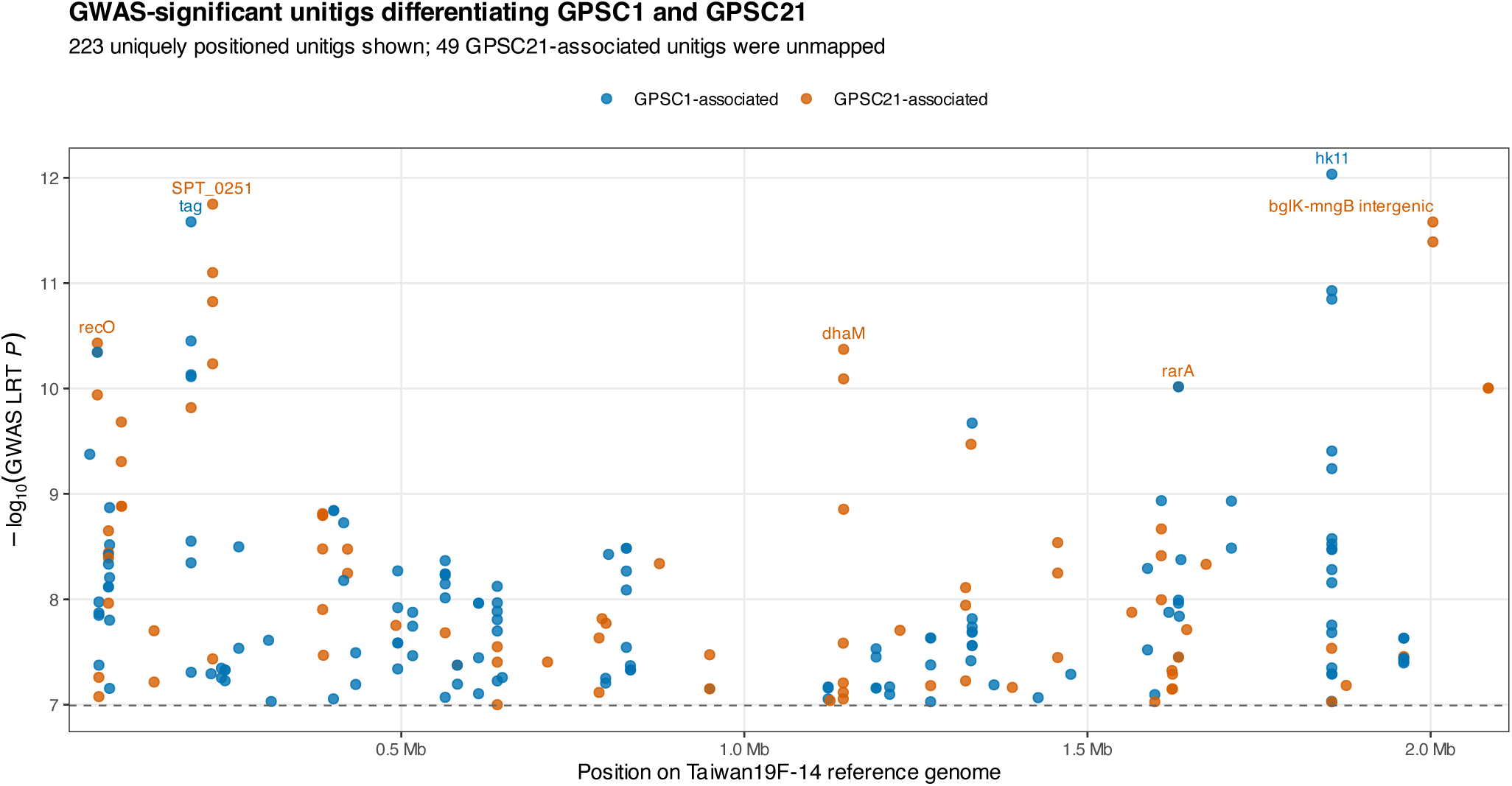
GWAS-significant unitigs differentiating GPSC1 and GPSC21. Among the 1,569 unitigs reaching significance in the invasion GWAS, we identified 272 that differentiated serotype 19F GPSC1 and GPSC21 isolates: 141 were GPSC1-associated and 131 were GPSC21-associated. Points show the 223 unitigs with a Taiwan19F-14 reference coordinate, positioned by genomic location and plotted against the negative base-10 logarithm of the original GWAS likelihood-ratio-test P value. Colours indicate the GPSC in which the unitig was enriched. The dashed horizontal line denotes the study-wide significance threshold, P = 0.05/490,378 = 1.01962 × 10^-7^, based on the number of unique unitig presence/absence patterns. All 141 GPSC1-associated unitigs mapped to Taiwan19F-14. Of the 131 GPSC21-associated unitigs, 82 mapped and are shown, whereas 49 did not map to the GPSC1 Taiwan19F-14 reference and are therefore absent from the positional panel. These 49 unmapped unitigs were rare in GPSC1 (frequency 0–4.2%) and common in GPSC21 (95.8–100%); their sequences, GWAS statistics, prevalence values and available locus annotations are retained in the accompanying unitig table. The highest-ranked rarA unitig was GPSC21-associated and unmapped; consequently, the rarA label is positioned using a lower-ranked mapped unitig from the same locus, whereas locus ranking in the manuscript is based on the highest-ranked unitig irrespective of mapping status.

